# Raster photostimulation of large-scale neural populations

**DOI:** 10.64898/2026.04.21.719951

**Authors:** Paul K. LaFosse, Daniel Flickinger, Georg Jaindl, Antonia Drinnenberg, Sverre Grødem, Kristian K. Lensjø, Charu Ramakrishnan, La’Akea Siverts, Hongkui Zeng, Bosiljka Tasic, Tanya L. Daigle, Marianne Fyhn, Karl Deisseroth, Carsen Stringer, Marius Pachitariu

## Abstract

Neural computations are implemented by distributed neural populations that typically span multiple brain areas. Simultaneous recording and photo-activation experiments can reveal the structure and logic of these neural computations, but such methods are typically limited to small subsets of neurons in restricted fields of view. Here we describe a new system called raster photostimulation for activating and recording thousands of neurons, over a short 300 ms time window and over a large 5 mm field-of-view on a two-photon mesoscope. The photo-activation is precisely matched to the neural recording configuration, as it uses the same optical path, although with a different laser that is independently gated. We demonstrate pixel-level precision, frame-by-frame mask updating, and single-frame photostimulation of thousands of neurons. While this method lacks the precise temporal control of alternative methods, it compensates with ease-of-use, spatial precision, cost of implementation and by pushing the limits on the number of near-simultaneously stimulated neurons.

## Introduction

Causal perturbations have played a key role in the study of neural computations, starting with lesion studies [1] and continuing with optogenetic approaches that have enabled precise targeting [2– 7]. When combined with one-photon light sources, optogenetics enables precise temporal and genetic targeting, although with diffuse spatial targeting due to the scattering of light in tissue [8–10]. Two-photon excitation allows for precise focal targeting despite scattering, but activating individual cells is difficult and requires advanced optical methods [11– 13]. The effort required for two-photon stimulation is well-rewarded, as it enables the study of many neural computations that span across molecular cell types, and thus cannot be targeted genetically. An example of such computation is orientation tuning in primary visual cortex (V1), where the feature selectivity of neurons to different orientations does not map onto molecular cell types, but rather onto input connectivity patterns [14]. Furthermore, different orientation-tuned neurons are interspersed in mouse V1, making diffuse spatial targeting impossible [15]. Thus, to perform causal perturbations in such circuits, two-photon optogenetics is required [16–22].

An additional challenge arises when studying higher-level computations involved in perception, cognition and action, which by their nature are distributed across multiple brain areas and across tens or hundreds of thousands of neurons. While two-photon optogenetics has been steadily improving, for example with various holographic methods [23–26], it remains mostly restricted to smaller fields of view (<1 mm2) and small neural populations of tens to hundreds of neurons [27, 16, 28–30]. Advanced methods to overcome these limitations are an active area of research and engineering, but they typically lead to substantial increases in the difficulty of implementing and maintaining the corresponding optical systems [31, 32]. This is made especially difficult by the requirement of simultaneously imaging the neurons [33], without which precise photostimulation maps cannot be obtained. Additional difficulty is added by the need to restrict photostimulation along the axial direction (*∼*10 *µ*m) to prevent off-target excitation above and below the targeted neuron.

Here we developed an alternative method that enables two-photon stimulation of thousands of neurons distributed over a wide region, with high spatial precision both horizontally and axially, while allowing for simultaneous large-scale neural recordings (Movie S1). To achieve these properties, the method trades off temporal precision, and requires the use of highly sensitive opsins such as ChRmine [16, 34, 20]. Below, we describe the procedure to achieve this by passing an additional red laser through the standard imaging optics and gating this laser at 20 MHz speeds in sync with the imaging configuration.

## Results

Our goal was to take advantage of the rapid scanning capabilities of standard resonant-galvo systems and use the same optical path for recording and photostimulation with two different lasers. We used a 920 nm green fiber laser for the recording, and a 1,035 nm “Monaco” red laser for photostimulation. Following pilot experiments, we used a 50Mhz repetition rate on the 1,035 nm laser with powers of up to 300mW at the sample. This power limit is also within range of most red fiber lasers, which should be substantially easier to operate than the Monaco we used here. We implemented the approach below on a large field-of-view two-photon mesoscope [35] as well as on a smaller FOV Thorlabs Bergamo 2 microscope. While both systems performed similarly, we we primarily describe our results on the two-photon mesoscope.

The main challenge to achieving effective raster photostimulation is the synchronization of signals between the gating signal sent to the red laser, and the raster scanning trajectories generated by the resonant-galvo system. We implemented this synchronization in ScanImage [36], together with a GUI system for manually drawing photostimulation targets. Each frame-pattern corresponded to a particular, binary waveform generated digitally at very high sampling rates, amplified and relayed to the AOM gate control on the red laser (see also [37] for an example of AOM control of the imaging laser). An additional binary signal was multiplexed with the waveform to specify which frames the waveform should be applied to.

In order to test the limits of this system, we started with a set of experiments in fluorescein slides. When applying a red mask pattern to a single resonant line of the recording, the pattern of readout fluorescence matched the photostimulation signal with some acquisition shot noise added (Figure 1a). Multiple resonant lines can be composed into a two-dimensional image patch with the usual method of bidirectional scanning (Figure 1b). Combined across multiple regions of interest, the system can generate pixel-accurate photostimulation masks across an entire field-of-view of over 5 mm on the mesoscope (Figure 1c). Applied to a fluorescein slide, this photostimulation mask generated photons in patterns that matched well the structure of the mask (Figure 1d), thus demonstrating the precision of the rasterized approach for photostimulation. In a different test, we used a fragment of the novel “Frankenstein” as the photostimulation mask, to further demonstrate the precision and scale of the photostimulation technique (Figure 1e,f).

**Figure 1:**
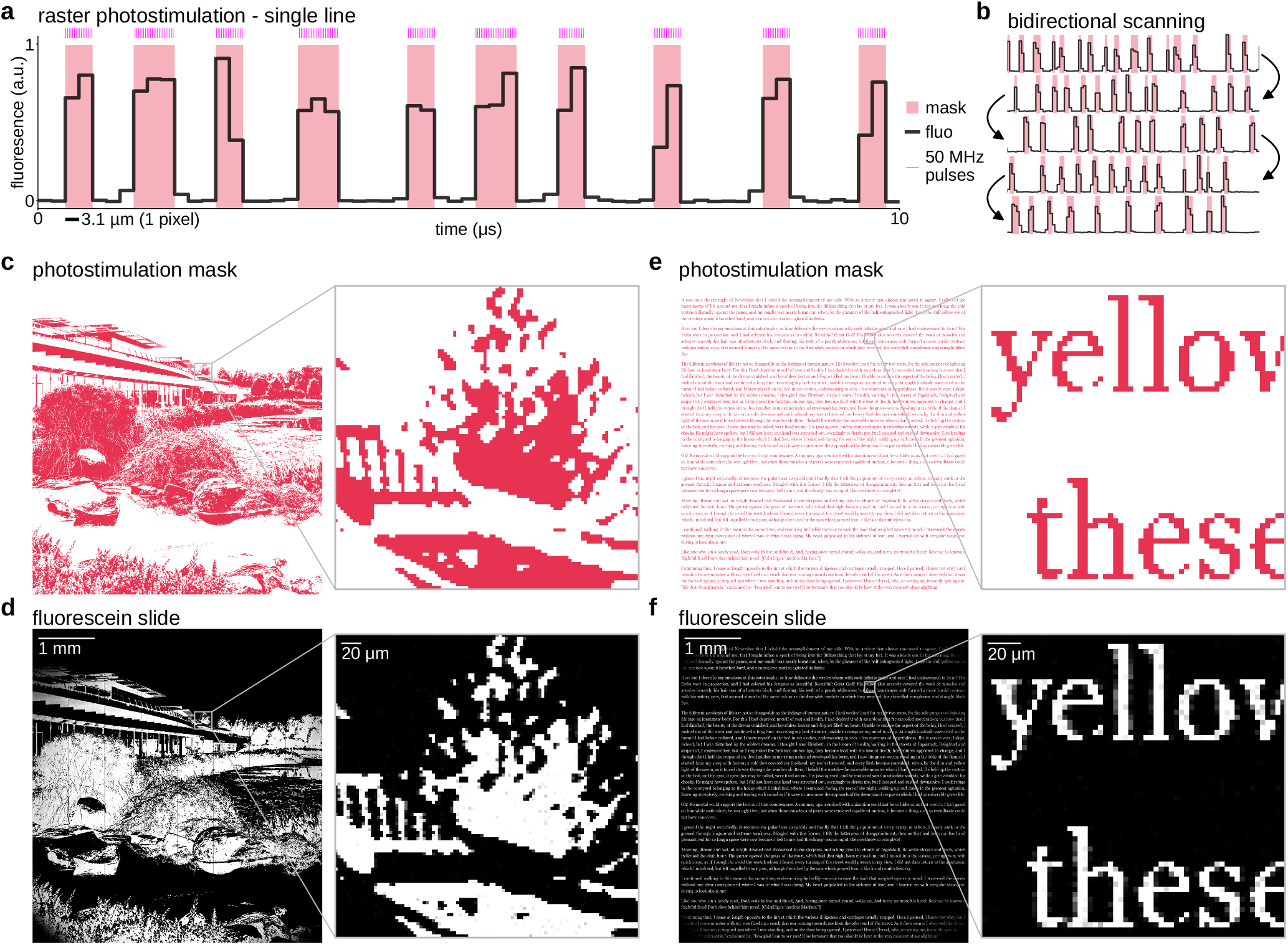
Sequential raster photostimulation. **a**, Portion of a single line during photostimulation with recorded fluorescence from a fluorescein slide, on the two-photon mesoscope. The mask indicates periods where the 50 Mhz stimulation laser was not gated. **b**, Portions of five consecutive lines illustrating bidirectional scanning with a different pattern in each line. **c**, Photostimulation mask that was designed for a 5.2 mm field of view. A zoomed-in patch is shown on the right. **d**, Recorded fluorescence from a fluorescein slide in response to the photostimulation mask in **c**. A zoomed in patch is shown on the right. **e-f**, Same as **c-d** for a different 5.2 mm pattern.

Next, we designed a protocol for recording and stimulating very large numbers of neurons. We first tested this protocol in fluorescein slides. We obtained photostimulation ROI masks from a previously-acquired recording (from [38]) in which over 32,000 cells were recorded in a single mesoscope plane (Figure 2a). Random subsets of 10,000 cells each were grouped into multiple photostimulation masks, with the cellular masks capturing the pixel-level details from Suite2p (Figure 2b) [39]. To demonstrate the application of these masks in quick alternation, we pre-computed a set of waveforms corresponding to each mask, and created a protocol for swapping masks in real time. The waveform swaps were performed in a 80 ms buffer period in-between consecutive frames. The recorded signals in fluorescein again matched very closely the masks that we applied (Figure 2c).

**Figure 2:**
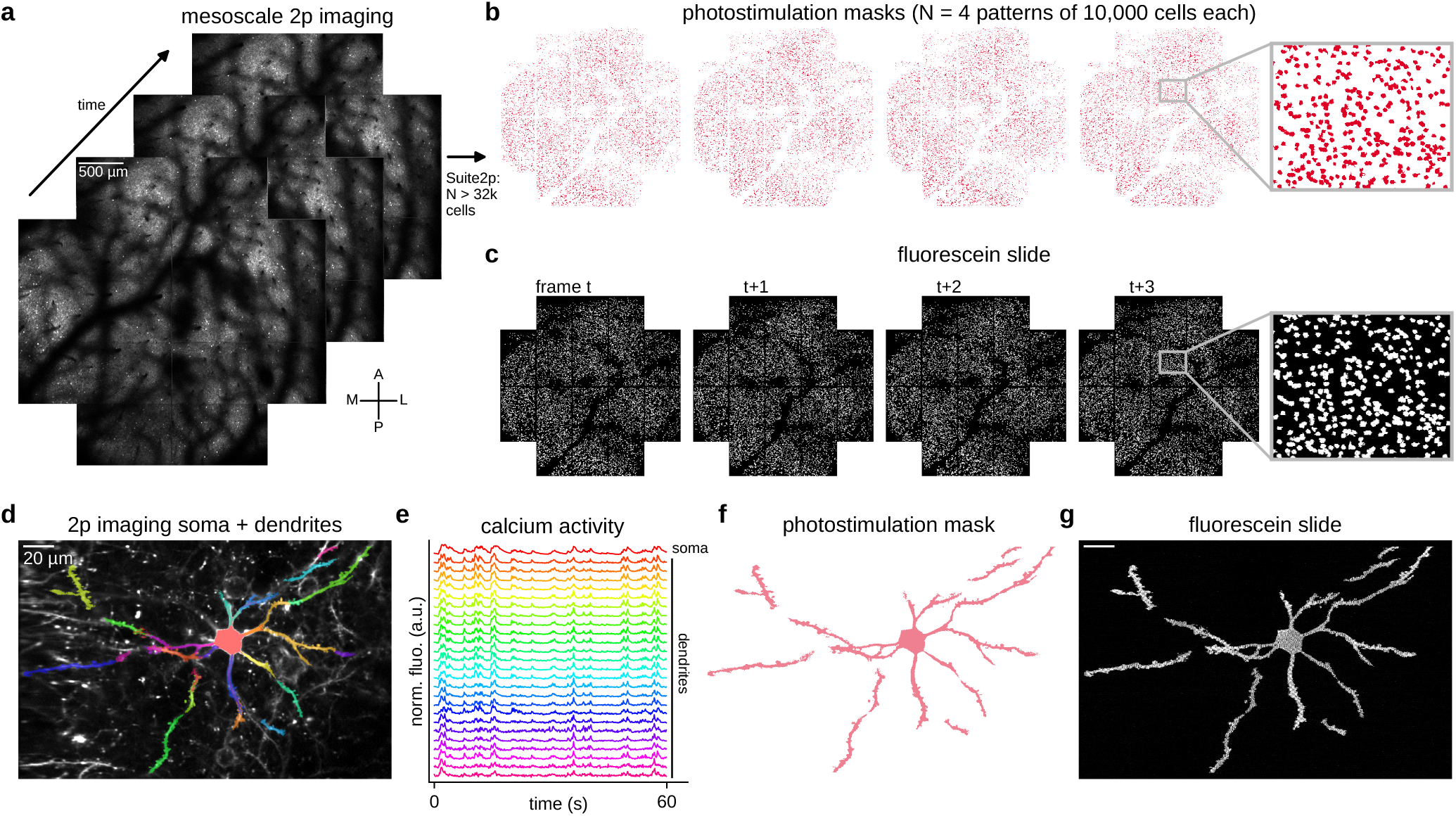
Designing photostimulation masks for *in vivo* experiments. **a**, Example frames from a large-scale two-photon recording on the mesoscope. Suite2p was applied to obtain over 32,000 cell masks. **b**, Multiple photostimulation masks created from random subsets of 10,000 cell masks from those identified in **a. c**, Two-photon recording of a fluorescein slide with frame-by-frame photostimulation with the photostimulation masks in **b**. The mask is changed on each frame. **d**, Mean image in gray and Suite2p masks in color from a zoomed-in two-photon recording from an experiment with sparse jGCaMP8s expression. **e**, Fluorescent traces recorded from the Suite2p masks in **d. f**, Photostimulation mask created from the active dendrite compartments and the soma of the cell in **d. g**, Single frame from a two-photon recording of a fluorescein slide during application of the photostimulation mask in **e**.

In addition to allowing for large-scale photostimulation, the raster photostimulation method allows for a high level of spatial precision. This can enable experiments where individual compartments on a neuron can be activated together or independently [40]. To demonstrate this, we recorded a neuron with several of its dendrites, in a sparse-expressing preparation (Figure 2d,e), and ran Suite2p on the recording. This produced multiple compartments, which we combined into a single mask (Figure 2e), and then applied to a fluorescein slide, again obtaining a faithful reproduction (Figure 2f) that demonstrates the precision of the raster photostimulation method.

Having demonstrated the spatial and temporal precision of raster photostimulation in fluorescein slides, we next turned to *in vivo* experiments. We used a triple virus injection strategy to express jGCaMP8s localized to the ribosomes, as well as ChRmine expressed in the same cells, with amplification provided by the tTA-TRE system [41, 34, 42]. This resulted in broad and relatively uniform expression of both constructs. After 4 weeks, we recorded neural activity from a mesoscopic field-of-view of 3.5 mm over primary and higher-order visual cortices at a frame rate of 3.18 Hz (Figure 3a). We ran this recording through Suite2p to obtain cell masks, and then selected a random subset of 1,000 neurons for photostimulation (Figure 3b). We applied this mask on individual frames of the recording, for a total of 100 repetitions or “trials”. After registering the raw recorded movie in Suite2p, we created pixel-level response maps by subtracting the pre-photostimulation average response from the frame immediately following the photostimulation frame (Figure 3c). The neural recording frame during photostimulation was acquired but not used, as it contained clear photostimulation artifacts. We also tested the method using Ai229 x Slc17a7-IRES2-Cre mice which express GCaMP6m and ChRmine in excitatory neurons and obtained similar results (Figure 3c, Movie S1), though in this experiment cells were targeted manually rather than obtained from Suite2p.

**Figure 3:**
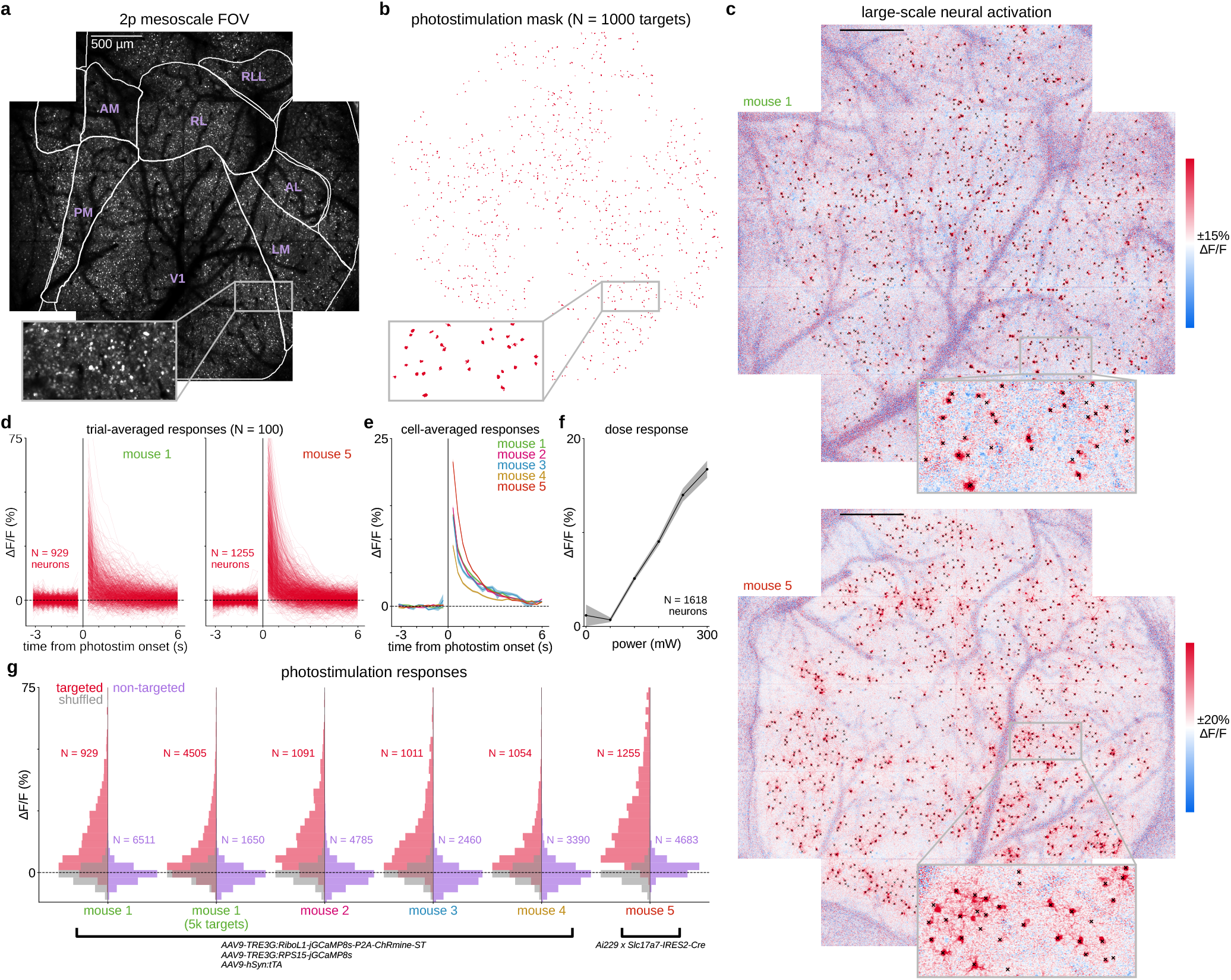
Large-scale “read and write” experiments *in vivo*. **a**, Example 3.5 mm field of view from the two-photon mesoscope. **b**, Example photostimulation mask targeting *∼*1,000 cells detected in a short recording without stimulation. **c**, Trial-averaged pixel responses to photostimulation with a fixed mask, for the duration of a single frame (N = 100 trials). Each pixel was baselined with its average value in the prior 10 frames before stimulation. Top: recording with viral expression of jGCaMP8s and ChRmine; Bottom: recording with transgenic expression of GCaMP6m and ChRmine. **d**, Trial-averaged response traces for the photostimulated cells in **c. e**, Trial- and neuron-averaged responses across mice (N = 5). Error bars represent s.e.m. across neurons (N = 929–1,255). **f**, Neural responses to photostimulation as a function of laser power. Error bars represent s.e.m. across neurons (N = 1,618). **g**, Histogram of response strengths for photostimulated cells compared to time-shuffled data from the same cells and compared to non-targeted cells.

At the single-cell level, a large fraction of targeted neurons across all mice had positive responses that could be detected on the frames following the photostimulation frame (Figure 3d,e). The responses were proportional to the amount of laser power used up to the tested limit of 300 mW. Note that while 300 mW was the power of the laser, it was blanked for approximately 99% of the frame, and each cell was only photostimulated for *∼*0.2 ms. While there was some variation in the strength of the responses, most targeted neurons responded substantially more than the non-targeted neurons (Figure 3g). The distribution of response strengths was almost entirely positive, compared to a shuffled response distribution obtained from non-photostimulation time points, which was symmetric around 0 (Figure 3g). The same photostimulation strategy could be scaled up to >4,000 neurons (Figure S1) and in principle to as many neurons as are available in the field of view.

We next characterized the robustness of raster photostimulation across various conditions. We first varied the point-spread function of the stimulation both by changing the beam size on the mesoscope, and by using a different two-photon microscope (Thorlabs Bergamo II) (Figure 4a-c). Across these configurations, we found robust activation of targeted neurons at constant stimulation power (Figure 4d). We also measured response differences in non-targeted neurons by changing PSF size on the mesoscope while keeping average power constant. In neurons <5 *µ*m from targeted neurons we found a small but significant increase in excitation using a larger PSF, but found no significant differences beyond 5 *µ*m (N = 5 mice; Figure 4e-g, Figure S2). While a smaller PSF may reduce “off-target excitation” (i.e. undesired, direct excitation), smaller responses in targeted neurons may also contribute to smaller responses in non-targeted neurons (Figure S3).

**Figure 4:**
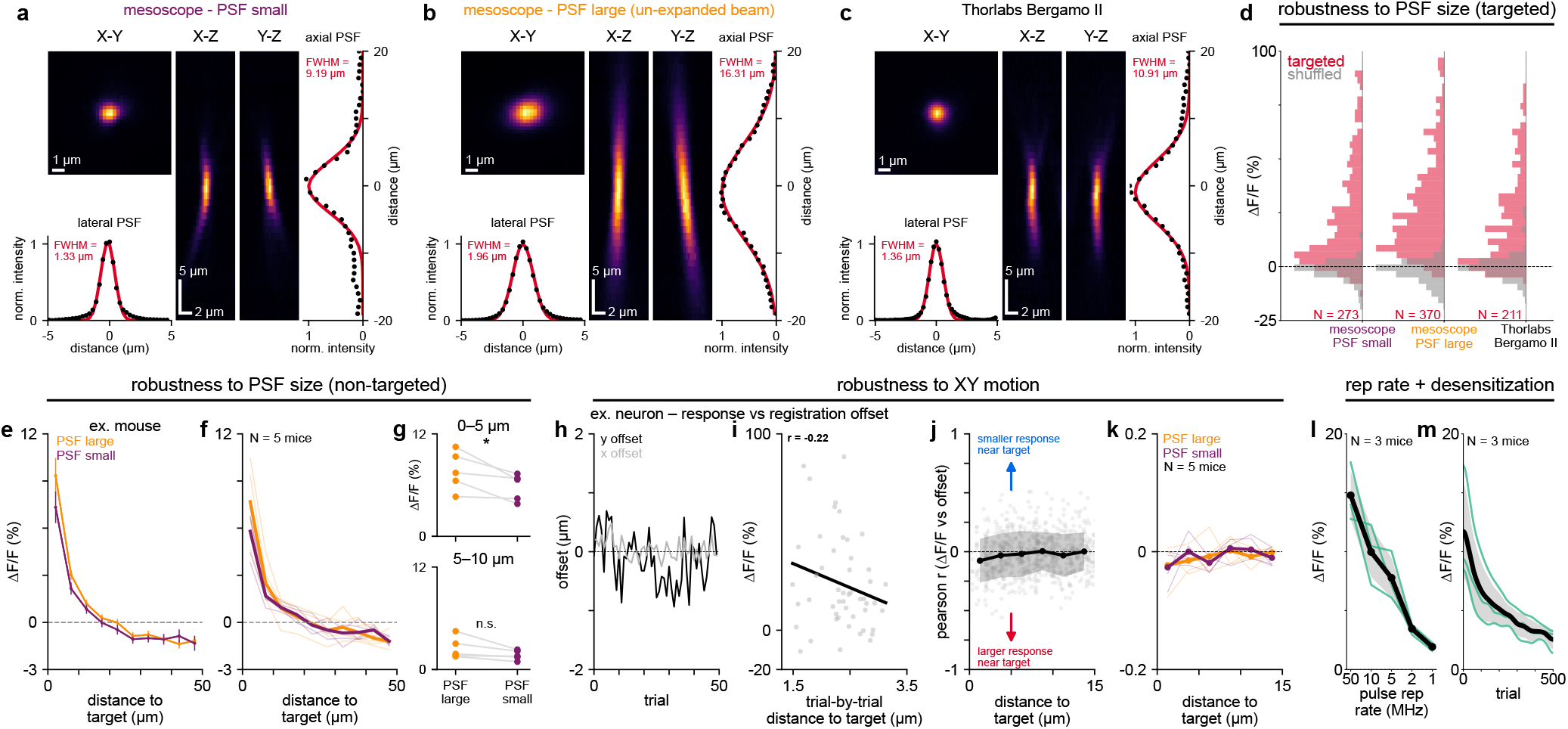
Characterization of raster photostimulation for *in vivo* use. **a**, Point-spread function (PSF) projections (2D and 1D) with gaussian fits for the mesoscope. **b**, Same as **a** for a mesoscope configuration with an un-expanded beam (used in the rest of the figures). **c**, Same as **a**, but using a Thorlabs Bergamo II microscope. **d**, Population responses to photostimulation (similar to Figure 3g). **e**, Example data of averaged responses as a function of distance to the nearest photostimulation target. **f**, Same as **e** across animals (thick line, mean; thin lines, individual mice). **g**, Paired average responses between large and small PSF sizes (N = 5; paired t-test, * p < 0.05). **h**, Non-rigid registration offsets for an example neuron. **i**, Relation between photostimulation response and distance to nearest target for example neuron across trials. **j**, Correlation across trials of response and distance to target for neurons <15 *µ*m away from a target in an example mouse. Black line and shaded region are mean ± standard deviation across neurons. **k**, Average curves like in **j** for all mice (thick lines, mean; thin lines, individual mice). **l**, Average photostimulation response in targeted neurons for different pulse repetition rates (mean ± standard deviation, N=3 mice). **m**, Average photostimulation response in targeted neurons across repetitions of photostimulation (mean ± standard deviation, N=3 mice).

We next characterized the robustness of photostimulation to motion of the field-of-view, such as that induced by animal movements in awake experiments. For each non-targeted neuron, we calculated the correlation of its response to the trial-by-trial distance from the nearest photostimulation mask (Figure 4h,i). If off-target excitation significantly impacts measured activity in nearby non-targeted neurons, then we might expect to find larger responses on trials where nearby neurons are closer to the photostimulation target due to motion (i.e. a negative correlation). However, across mice (N = 5) we found no effect for nearby neurons (Figure 4j,k), demonstrating that raster photostimulation is robust to small amounts of in-plane motion.

Finally, we examined the influence of laser pulse repetition rate and repeated stimulation on raster photostimulation. While keeping total output power constant (210 mW), we found that higher pulse rates led to larger neural responses (Figure 4l), likely due to more uniform coverage of pulses over the neuron area. Of note, using 1 MHz pulses (with high peak pulse power) on occasion led to spreading depression-like activity across the imaging FOV [43]. At our highest available pulse rate (50 MHz), we next tested the robustness of responses to repeated raster photostimulation, as prolonged imaging and photostimulation can result in opsin desensitization [18, 44]. We observed a modest reduction of 25 % in target responses after 50 trials (300 mW stimulation), implying that the method can be used effectively in typical experiments. However, after continued stimulation of the same cells for 500 trials, we observed a marked reduction in responses by 78% (Figure 4m).

## Discussion

Here we demonstrated a method for two-photon optogenetics that can be implemented on nearly any raster-scanning two-photon microscope, at relatively low cost. The method ties photostimulation parameters to the imaging parameters, which has both pros and cons. On the positive side, it enables photostimulation of as many neurons as can be recorded simultaneously, in a large field of view, while using point-spread functions as small as the imaging system allows, thus minimizing off-target stimulation. While we have primarily shown the method implemented on a large 25 mm^2^ field-of-view mesoscope, we have also implemented it on a Thorlabs Bergamo II microscope, which can record fields of view up to 4 mm^2^ [39]. Multiplane imaging can be used in this case to target raster photostimulation to more neurons across multiple planes. Since the Bergamo II can be used to record over 100,000 neurons [39], it follows that it can also be used to raster photostimulate over 100,000 neurons, though we have not conducted such experiments yet.

On the negative side, the temporal precision of raster photostimulation is much reduced compared to holographic photostimulation, and the order of temporal activation is tied to the order of raster scanning the neurons. Similar to raster imaging, there is some flexibility in raster photostimulation to trade off temporal accuracy against the number of neurons recorded: one may image a large field-of-view at slow frame rates, or a small field-of-view at high frame rates. Since all recorded neurons can be photo-activated within the period of a single frame, higher frame rates result in more temporal precision. Alternatively, new types of microscopes may be used that can inherently image faster [45], which would directly translate to faster and more temporally precise photostimulation. Note that on some holographic systems and microscopes, including the 2p-RAM, it is possible to add a galvo-galvo relay into the photostimulation path in order to temporally multiplex patterns and increase the number of neurons near-simultaneously photostimulated [31, 32]. This, however, introduces similar temporal tradeoffs as raster photostimulation and is generally harder to implement and maintain. As an additional challenge to raster photostimulation, there is a non-uniform distribution of laser power across a raster line due to changes in velocity of the resonant scanner, up to a factor of *∼*3 between the center and edge of the line. This can be corrected for by modulating laser power across the raster line.

Our method is complementary to temporally-precise holography. While holography can answer questions about the precise order of activation of neurons [28, 46], raster photostimulation can answer questions about the scaling of photo-activation with large numbers of neurons. Recent studies have probed the influence on behavior of stimulating tens of neurons [16, 27, 47–54], which could be expanded to thousands of neurons with raster photostimulation. Beyond biasing perceptual decisions and motor commands, large-scale photostimulation can also be used to investigate connectivity patterns and circuit motifs within and across brain areas, both at the level of single neurons and at the level of subpopulations of coordinated neurons [55–59, 30, 60–63]. It may also be used to interrogate cortical plasticity rules through repeated photostimulation [64–66] or pre/post connectivity measurements [67], at a larger scale than previously explored.

## Acknowledgments

This research was funded by the Howard Hughes Medical Institute at the Janelia Research Campus. From the Vivarium, we thank Jim Cox, Crystall Lopez, Anne Kuzspit, Miriam Rose, Amanda Minisi, Alexa Gracias, Gillian Harris, Sarah Lindo, and their respective teams for animal breeding, husbandry, and surgeries. From JeT, we thank Vasily Goncharov, Alex Sohn, Tobias Goulet, and Steven Sawtelle for help with rig maintenance and upgrades. From Molecular Genetics and Viral Tools we thank Kym Delventhal, Alex Ludlow, and Hyun Ah Yi. From MBF Bioscience we thank Boris Djiguemde and Samuel Ventura for ScanImage support.

## Author information

## Author contributions

P.K.L., C.S., and M.P. designed the study. D.F. developed and maintained the microscope and its upgrades. G.J., M.P., and P.K.L. conceptualized and developed the raster photostimulation control software. S.G., K.K.L., and M.F. developed viral tools for ribosomal GCaMP expression. A.D., C.R., L.S., H.Z., B.T., T.D., and K.D. developed transgenic mice for ChRmine expression. P.K.L. performed photostimulation experiments. P.K.L., with input from C.S. and M.P., performed data analysis. P.K.L., C.S., and M.P. wrote the manuscript with input from all authors.

## Methods

We used Python 3 [68] for data analysis, using numpy, scipy, pytorch, numba, scikit-learn, tifffile, and ScanImageTiffReader [69–75]. Figures were made using matplotlib and jupyter-notebook [76, 77].

### Data acquisition

#### Animals

All experimental procedures were performed in accordance with the IACUC at HHMI Janelia Research Campus. For *in vivo* experiments, we used both transgenic and viral strategies to co-express GCaMP calcium indicators and the opsin ChRmine in cortical neurons of mice. We recorded from one Ai229 x Slc17a7-IRES2-Cre mouse (RRID:MGI:7482008, RRID:IMSR_JAX:037512), one Ai228 x Cux2-CreERT2 mouse (RRID:MGI:7482006, RRID:MGI:5011472) [20], one TetO-GCaMP6s x CaMK2a-tTA mouse (RRID:IMSR_JAX:024742, RRID:IMSR_JAX:003010) [78], as well as eight VGAT-Cre x Ai14 mice (RRID:IMSR_JAX:028862, RRID:IMSR_JAX:007914). All mice were female and ranged from 7 to 12 months of age. Mice were housed with a reverse light cycle and paired with a sibling before and after surgery. Holding rooms were maintained at a temperature of 70° F ± 2° F and humidity of 50% rH ± 20%.

#### Surgical procedures

Surgical methods are described in detail in previous work [79]. In short, craniotomies were performed in adult mice (between ages of 3 and 7 months) anesthetized with 3% Isoflurane (1.5-2% during procedure) to implant a 4+5 mm diameter circular double window over visual cortex (4 mm window inserted to fit craniotomy and 5 mm window fitted over the edge of the bone). During the procedure, Marcaine (*≤*8 mg/kg) was initially injected subcutaneously under the incision area, and followed by warmed fluids + 5% dextrose and Buprenorphine (0.1 mg/kg) administered subcutaneously along with Dexamethasone (7 mg/kg) via intramuscular route. A custom stainless steel headbar was positioned around the craniotomy site prior to drilling and the full implant was secured in place using UV cured resin cement (Calibra Universal). Following the procedure and for 2 days after, mice were administered Ketoprofen (5 mg/kg) subcutaneously and monitored for pain or distress.

In the VGaT x Ai14 mice, viral injections were performed during the procedure prior to implanting the window. Three viruses were mixed together in PBS: AAV9-TRE3G:RiboL1-jGCaMP8s-P2A-ChRmine-ST (final titer: 4.25 *\**10^12^ GC/mL), AAV9-TRE3G:RPS15-jGCaMP8s (final titer: 1.06 *\**10^13^ GC/mL), and AAV9-hSyn:tTA (final titer: 6.98 *\**10^12^ GC/mL for mice 1-4, and 2.79 *\**10^13^ GC/mL for mice 6-9) (RiboL1-jGCaMP8s described in [41]). Injections of 100 nL volume were administered at two depths (300 and 500 *µ*m) at each of 3 sites spanning the exposed cortical surface within the craniotomy (6 total injections per mouse).

#### Imaging acquisition

During recordings, head-fixed mice were free to run on a treadmill, having been acclimatized to the treadmill prior to recordings.

We used a two-photon mesoscope [35] for recording photostimulation-induced fluorescence measurements and for recording neural activity. We also used a Thorlabs Bergamo II with a 16x Nikon objective for validation on a standard two-photon microscope (Figure 4). ScanImage acquisition software was used for data collection [36]. For neural recordings, online motion correction was used to correct drift in both *z* and *xy* axes [79]. A 920 nm fiber laser (30 MHz pulse rate; FemtoFiber Ultra, Toptica) was used to image neural activity at 30–48 mW of power delivered to the sample.

Mesoscale recordings (Figure 2a, Figure 3) were performed across a 3.5 mm diameter field-of-view (1.3 and 2.0 *µ*m/pixel in x and y dimensions, respectively) at 3.18 Hz. A zoomed-in recording of neurons (Figure 2d) was taken using the mesoscope with six ROIs of size 190 × 130 *µ*m (∼0.3 *µ*m/pixel) each at 7.5 Hz - one ROI was used for Figure 2d. For comparisons of different point-spread functions (PSFs) and systems (Figure 4), zoomed-in recordings on the mesoscope were performed using two ROIs spanning a square FOV of size 1.3 × 1.3 *µ*m and the Thorlabs Bergamo II microscope with a single ROI of size 1.3 × 1.3 *µ*m.

#### Raster photostimulation

A 1,035 nm laser (50 MHz pulse rate, except in Figure 4l, and 277 fs pulse width; Monaco, Coherent, Inc.) was coupled into the optical path of the imaging laser for raster photostimulation. Photostimulation power ranged from 70–300 mW (Figure 3 and Figure 4). For *in vivo* experiments, cells only received photostimulation laser pulses over a brief window of time, roughly proportional to the fraction of total scanning area occupied by a single photostimulation mask. For scanning a full mesoscopic field-of-view (Figure 3a) at 3.18 Hz with a 2440 x 1716 pixel resolution and a cell-defined photostimulation mask of 43 pixels on average, we estimate a given cell receives

photostimulation pulses for *∼*3.2 *µ*s over an entire imaging frame, where *∼*1 *µ*J of energy is delivered using a 300 mW laser output. Due to the line-by-line scanning of the laser, a cell was typically stimulated with 4-6 bursts of pulses over an interval of *∼*0.2 ms. The number of laser pulses per pixel in a raster line will be greater for pixels at the edge than at the center as the resonant scanner slows to turn around, which impacts the amount of energy delivered to the sample. For the 5.2 mm field-of-view (Figure 1), the number of pulses per pixel ranged from 5.9 to 18.5 with a mean of*∼*8 pulses per pixel. For the 3.5 mm field-of-view (Figure 3), the number of pulses per pixel ranged from 2.2 to 8.1 with a mean of ∼3 pulses per pixel. Timing of the photostimulation was controlled using an external gate for an acousto-optic modulator (AOM) on the laser. The gating signal was composed of a 20 MHz TTL that conveyed the waveform produced by the photostimulation mask uploaded to ScanImage. A TTL logic gate (Pulse Research Lab) was used to filter the pattern waveforms to the AOM to only allow emission of laser pulses during specified frames during an acquisition.

For *in vivo* photostimulation experiments, to define photostimulation masks based on neuron morphology (Figure 3b), we first passively recorded calcium activity for 15 minutes. We then quickly processed the recording in Suite2p (“Ca-timescale” = 0.25 mice 1–4 and 6–9, 0.7 mouse 5, otherwise default parameters) to extract cell masks, from which *∼*1,000 (or 5,000; Figure S1) cell masks were randomly chosen to construct a single binary image. This image was uploaded to a customized module in ScanImage to compute the appropriate waveform for timing the photostimulation pulses to the selected neurons during scanning. For the zoomed-in field-of-view (Figure 2d), we used Suite2p with the same parameters as above but “Highpass neuropil” = 150 to generate masks of dendritic compartments and the soma, and constructed the full photostimulation pattern via manual selection of only the masks belonging to the cell of interest based on visual inspection of calcium activity across the components (Figure 2d-g). We used Suite2p cell masks from a recording from [38] for Figure 2b.

#### Photostimulation PSF

Fluorescent beads (1 *µ*m; Fluoresbrite YG Microspheres, Polysciences, Inc.) suspended in agarose were used to measure PSFs of the photostimulation laser under different conditions and across microscopes [80, 81]. A volumetric stack of the beads was first acquired and then analyzed to identify bead locations based on peak intensity and to measure the intensity profile across individual beads. For a given bead, intensity was collapsed to a one-dimensional axial profile (mean over both lateral axes, X and Y) and a lateral profile (the average of two profiles, one per lateral axis, each computed as the mean over Z and the orthogonal lateral axis). The full-width at half-maximum of a Gaussian fit to these profiles defined the axial and lateral PSFs. A beam expander was added to the light path of the two-photon mesoscope to decrease the lateral and axial PSF for response comparisons (Figure 4). All other photostimulation experiments were performed without the beam expander in the light path.

## Data analysis

### Photostimulation response quantification

We ran Suite2p on the photostimulation recording period to detect neurons and extract fluorescence for analysis, excluding frames with photostimulation. We used the fluorescent trace outputs *F* to compute Δ*F/F*_0_ for each neuron. *F*_0_ was computed as the average baseline activity prior to photostimulation periods (10 frames at 3.18 Hz, or 3.1 seconds; 60 frames at 17.46 Hz, or 3.4 seconds) across all repetitions (N = 30–100 repetitions, or trials). For time courses of responses (Figure 3d), Δ*F/F*_0_ activity was averaged across all photostimulation repetitions. Singular response values (Figure 3f,g) were defined as the trial-averaged Δ*F/F*_0_ activity from the first frame after the photostimulation frame (Figure 3, Figure 4d) or the average across all frames of photostimulation for non-targeted neurons (5 frames at 17.46 Hz, or 286 ms; Figure 4e–k). For quantification of desensitization across repetitions (Figure 4m), we measured the incremental increase in Δ*F/F*_0_ at each trial by subtracting the average preceding activity (10 frames at 3.18 Hz, or 3.1 seconds) from the response to photostimulation.

To identify neurons as either targeted or non-targeted, we computed the percent overlap of the photostimulation pattern mask to the masks of the recorded cells. For mice 1–4 and 6–9, any cell whose mask overlapped with a photostimulation mask *≥*20% was considered targeted. For mouse 5, which instead used larger squares (15 *µ*m width) over targeted cells instead of photostimulation masks defined by the cell morphology, we set an overlap threshold of *≥*80% to identify the cells primarily under the target mask. Non-targeted cells in all cases were defined as having 0% overlap with any part of the photostimulation mask.

For visualization of responses across the field-of-view (Figure 3c), Suite2p-registered and background subtracted images were used to compute trial-averaged Δ*F/F*_0_ values at each pixel using the same time window for *F*_0_ as described above and averaging the first 3 frames (1 second) of activity following photostimulation to compute *F*.

Distance to the nearest target was computed as the length of the vector between the closest pair of pixels in a cell’s Suite2p mask and the photostimulation mask. This vector’s X and Y components summed three terms: the raw pixel distance between the cell mask and photostimulation mask; the static offset of the photostimulation pattern, calculated by cross-correlating the Suite2p registration reference images (“refImg”) from the recordings for generating the pattern versus measuring photostimulation responses; and non-rigid registration offsets from Suite2p, taken either as each cell’s mean position across all photostimulation frames (Figure 4e–g, j, k) or its per-trial mean position (Figure 4i). Non-rigid offsets were computed per pixel within each cell’s mask and averaged at each time point (for details, see [39]).

---

**S1:**
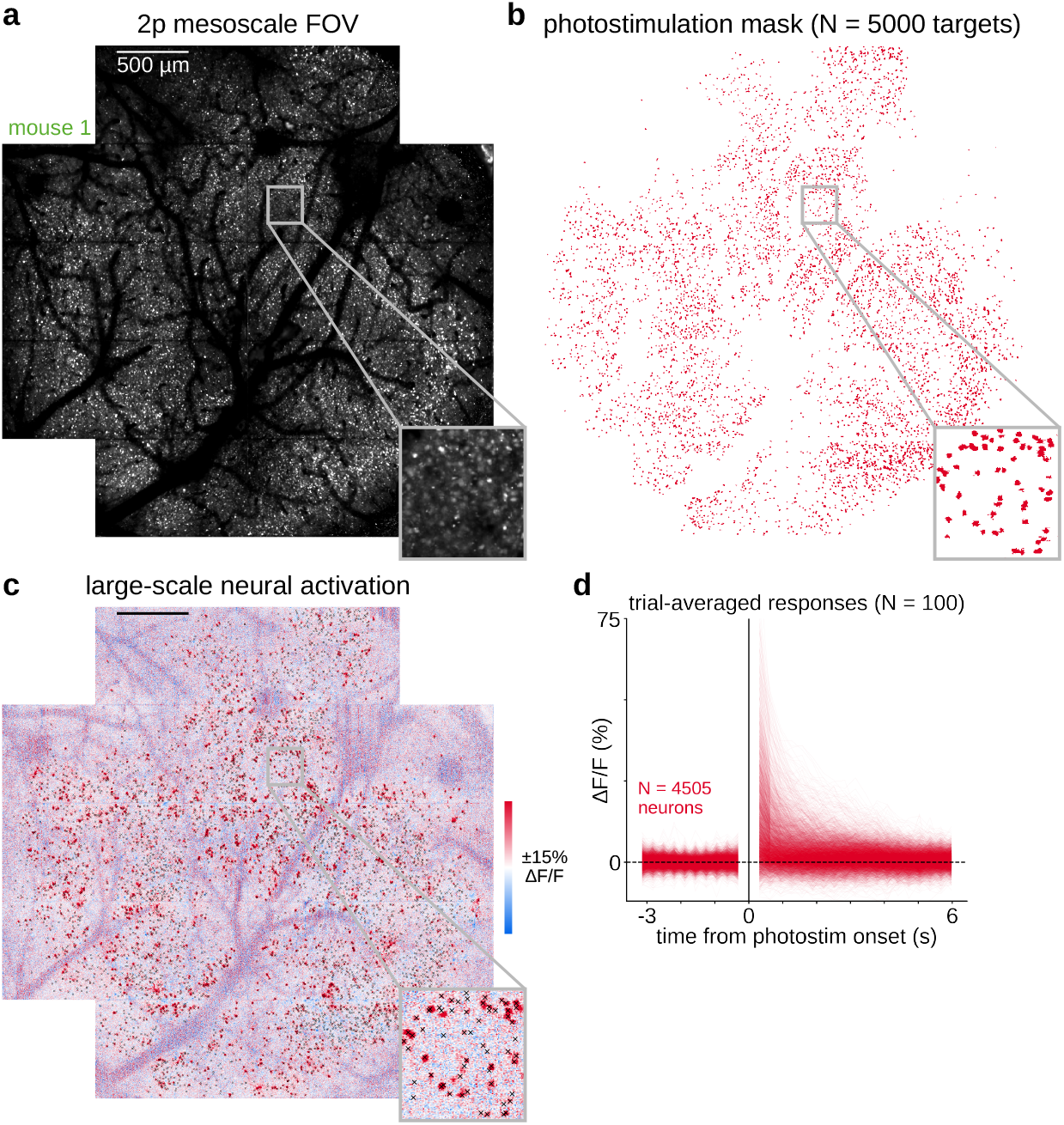
Photostimulation of thousands of neurons *in vivo*. **a**, Example 3.5 mm field of view. **b**, Photostimulation mask targeting *∼*5,000 cells. **c**, Trial-averaged (N = 100) pixel responses to photostimulation of the mask. **d**, Trial-averaged response traces of cells photostimulated.

**S2:**
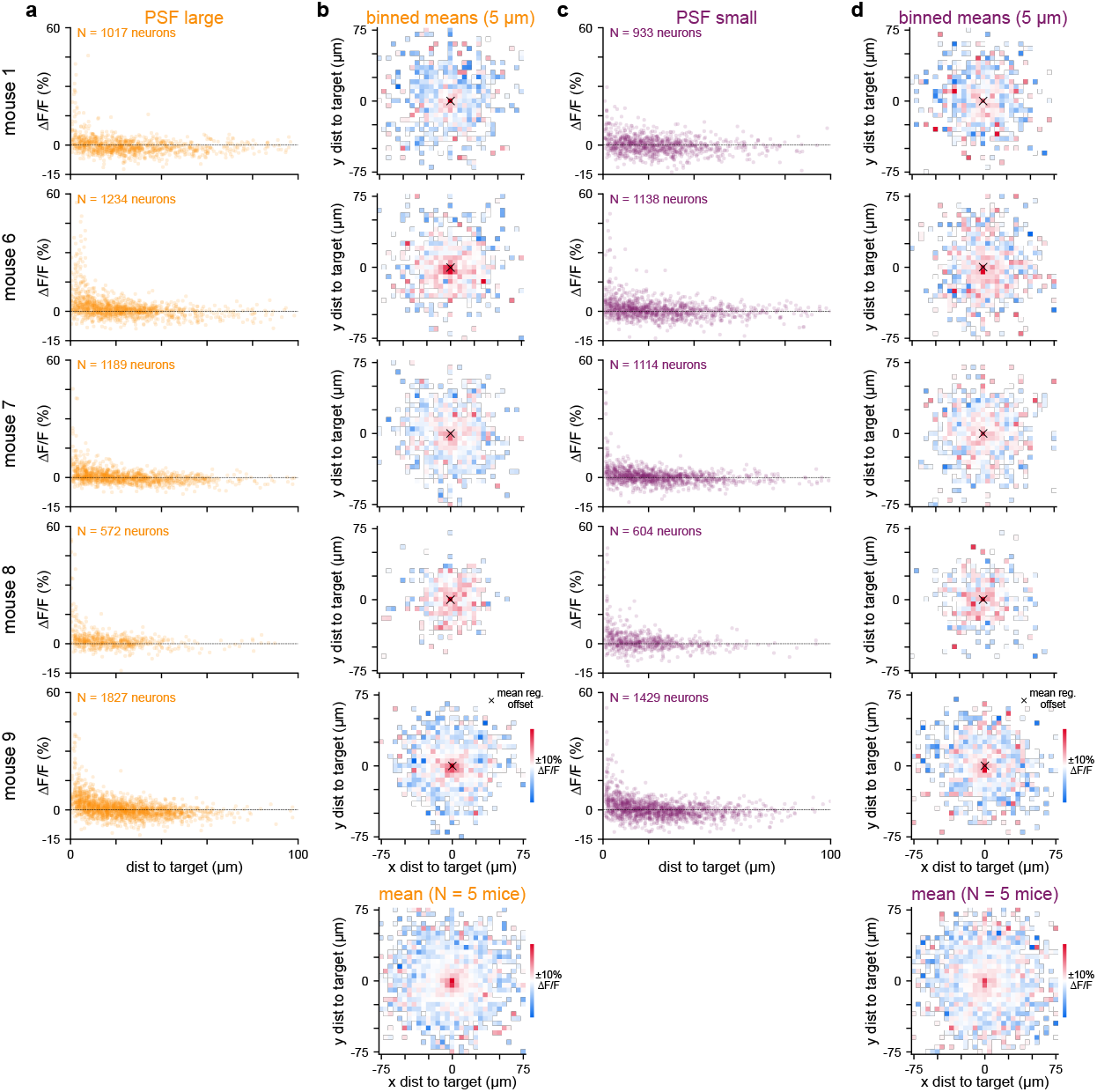
Photostimulation-evoked responses in non-targeted neurons as a function of distance to targeted neurons. **a**, Non-targeted neuron data for all mice using a large PSF. Trial-averaged responses across photostimulation frames (N = 50 trials, N = 5 frames at 17.46 Hz, or 286 ms) as a function of distance to the nearest pixel of the photostimulation mask. **b**, Responses averaged in 5 *µ*m bins and split into X and Y distance components. Bottom, mean binned responses across all mice. **c–d**, same as **a–b**, but for a small PSF.

**S3:**
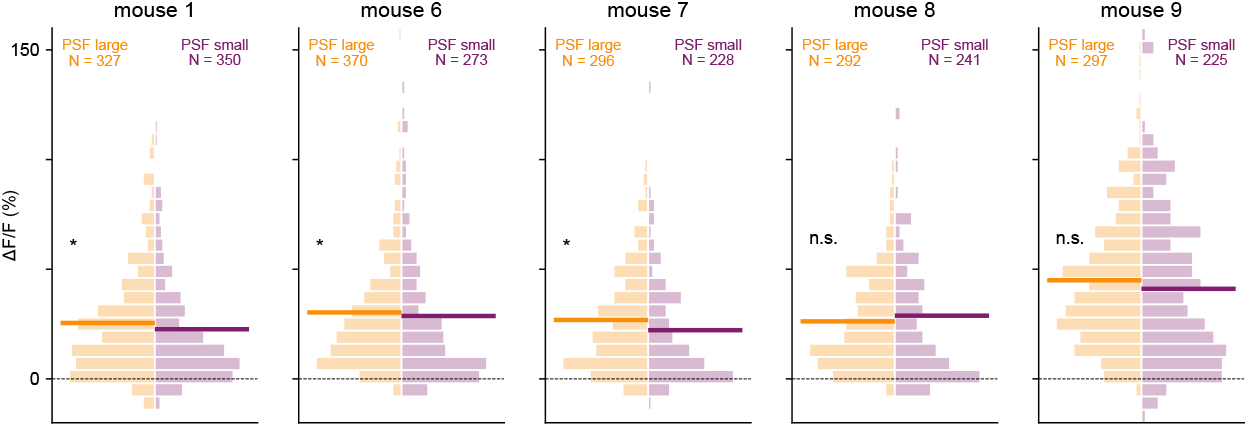
Photostimulation responses in targeted neurons for varied PSF size. Histograms of response strengths across targeted neurons at two different PSF sizes (N = 5 mice). Solid lines represent the mean response. Changes in responses due to different PSF quantified using a Mann-Whitney U test (* p < 0.05).

**Movie S1**. Movie of average pixel responses to photostimulation (N = 100 trials), baseline subtracted by average pre-stimulation activity. 10 frames before and 13 frames after stimulation (3.18 Hz frame rate) are shown at 2x real-time. Available at https://www.youtube.com/watch?v=TIlc5XtmxsQ.

